# Large deletion variants in a *Plasmodium falciparum* ookinete protein gene and associations with different endemic populations and mosquito vectors

**DOI:** 10.64898/2026.08.03.742444

**Authors:** Samuel Palmer, Balotin Fogang, Lindsay B. Stewart, Kate Chasseaud, Mojca Kristan, Thomas G. Anthony, Antoine Claessens, David J. Conway

## Abstract

The *Plasmodium falciparum PIMMS43* gene encodes an ookinete protein that is important for mosquito infection. Here, large indel variants within the coding sequence are identified, and frequencies in natural infections of humans and mosquitoes investigated. Comparing long-read genome sequences in a small panel of *P. falciparum* strains and related species revealed a 150 bp deletion in the central part of the gene in several strains including the standard 3D7 reference genome. Mapping of short-read genome sequence data from 524 *P. falciparum* infections from nine different countries to full-length *PIMMS43* showed the 150 bp deletion and an alternative 90 bp deletion to be common structural variants. Across African populations, the 150 bp deletion had a mean allele frequency of 32%, and the 90 bp deletion a mean frequency of 12%, with significant geographical variation. Targeted genotyping of 30 oocysts from naturally infected mosquitoes in Tanzania by nested PCR showed overall deletion variant frequencies similar to those seen in human infections in the same country. Although sample size was limited, a difference in the variant frequencies in oocysts from *Anopheles gambiae* and *Anopheles funestus* suggests potential vector-specific selection. The findings highlight the importance of surveying structural genomic variation and its potential role in parasite adaptation.

**Data summary:** The long-read genome sequence data sources are listed in the Supplementary Information (Supplementary Table 1). The short-read genome sequence data from human infections were extracted as a selected subset of those in the Pf7 release of global data from the MalariaGEN Consortium as described in the Methods. The SNP and indel genotypes derived from each sample of parasites cultured in the laboratory and from natural mosquito infections are given in the Supplementary Information (Supplementary Tables 1 and 3, Supplementary Figure 1).

**Impact statement:** This study identifies large indel polymorphisms in the malaria parasite transmission-stage gene *PIMMS43*, reflecting substantial structural variation previously overlooked by genome-wide SNP-focused analyses. Using long- and short-read genomic data across diverse samples followed by targeted genotyping, we show that deletion variants are frequent and geographically structured, with initial evidence consistent with potential vector-specific selection. This highlights the importance of incorporating structural variation into population genomic surveillance and studies of parasite adaptation.

## Introduction

Extensive studies have been performed on genomic variation in the major human malaria parasite *Plasmodium falciparum*, although most description of polymorphisms has involved mapping of sequences to a single reference genome ^1,2^. A single reference is not expected to contain all of the genomic template features needed to identify variant features common among other individuals of the species. Long-read genome assemblies of laboratory strains and short-term cultured clinical isolates ^3^, including strains used in experimental vaccine experiments ^4^, provide information that can be examined to identify such variants. Malaria parasite genomes are highly AT-rich, which affects mutation patterns ^5^ and leads to many small indel variants ^6^. Large indels have been less systematically characterised, although they are likely to contribute to parasite adaptation.

Studies on malaria parasites have mostly focused on asexual blood stages which cause disease in humans and can also be grown in culture. However, it is sexual stages that infect mosquitoes, in which they face a range of innate immune defences ^7^. The single-locus *P. falciparum* gene *PIMMS43* (*Plasmodium* Infection of the Mosquito Midgut Screen 43) encodes a protein functioning in infection of mosquitoes ^8^. The orthologue in the rodent malaria parasite *P. berghei* was termed *POS8* or *PSOP25* in previous independent studies ^*9,10*^, being highly transcribed in the motile ookinete stage that invades the mosquito midgut ^11^, and the protein detected in ookinetes as well as sporozoites. Experimental gene knockout analyses indicate that the PIMMS43 protein may function to prevent ookinete clearance by mosquito innate immune defences ^8^. The protein may also be required for the growth of sporozoites, as inactivation of mosquito complement-like responses allows *PIMMS43*-knockout parasites to undergo oocyst formation but not sporozoite maturation.

In the *P. falciparum* reference strain 3D7, *PIMMS43* is a single copy gene on chromosome 6 (locus PF3D7_0620000) without any intron, encoding a protein of 505 amino acids in length with an N-terminal signal peptide, and a C-terminal transmembrane domain with conserved cysteine residues ^12^. A preliminary analysis of single nucleotide polymorphisms (SNPs) in the *PIMMS43* coding sequence suggested that some of the variation may be differentiated among populations in Africa ^8^. The transmission-blocking potential of the product justifies further investigation of variation in *P. falciparum* populations.

Here, common deletion variants are discovered in the *PIMMS43* gene, by first examining available long-read sequences and then mapping short-read sequences of a wider set of samples onto a reference sequence with an intact non-deletion allele. This enabled the frequency of variants to be determined in widely distributed population samples covering most of the endemic range of *P. falciparum*, revealing significant geographic variation. Targeted genotyping was then used for analysing parasites in wild-caught mosquitoes. As each parasite oocyst contains up to a few thousand sporozoite progeny of a single zygote, DNA isolated from individual oocysts may be analysed to derive genotypes of the diploid zygotic phase ^13-16^. The *PIMMS43* deletion variant frequencies were determined in a population sample of oocysts from natural infections of *Anopheles gambiae* and *An. funestus* mosquitoes collected in Tanzania. This enabled exploration of potential differences in composition of oocysts in different vector species, and comparison with data from human infections.

## Methods

### *P. falciparum* DNA samples from cultured parasites and naturally-infected mosquitoes

Genomic DNA was analysed from a panel of 14 *P. falciparum* cultured laboratory strains of diverse origins (3D7, Dd2, T9/96, T9/102, RO33, K1, MAD20, Palo Alto, FCR-3, HB3, Wellcome, FCC-2, 7G8 and D6). This panel represented a broad range of the diversity within *P. falciparum*, as previously described in surveys of gene copy number variation and other gene sequence polymorphisms ^17,18^.

*PIMMS43* variation in mosquito infections was analysed by PCR of DNA from 30 *P. falciparum* oocysts isolated from midguts of 25 naturally infected female *Anopheles* mosquitoes previously collected from houses in Muheza, Tanzania ^15^. Of these, 11 infected mosquitoes were *An. gambiae* sensu stricto (from which 14 oocysts were genotyped), and 14 infected mosquitoes were *An. funestus* sensu lato (from which 16 oocysts were genotyped). Most of the oocysts were a subset of those in a previous study of other polymorphisms ^15^, with limited material remaining. Given the small amount of DNA available from each oocyst, whole genome amplification was performed to enrich *P. falciparum* template abundance using a pool of primers as previously described ^19^, prior to performing targeted PCR on the *PIMMS43* gene.

### *PIMMS43* sequence alignments to discover and genotype indels and SNPs

Investigation of *PIMMS43* variation was first done by comparing sequences of 19 *P. falciparum* strains, using genomes assembled after PacBio long-read sequencing in two previous studies ^3,4^, together with *PIMMS43* orthologues from related species in the *Laverania* sub-genus, *P. reichenowi* and *P. praefalciparum* ^20^. Sequences were retrieved using the PlasmoDB browser (www.plasmodb.org) ^21^, and individual sequence IDs are listed in Supplementary Table 1. A multiple sequence alignment was performed on all sequences in FASTA format using Clustal Omega v1.2.4 (on www.ebi.ac.uk/jdispatcher/msa/clustalo) ^22^, and alignments were visualised with the NCBI Multiple Sequence Alignment Viewer v 1.25.0 (on www.ncbi.nlm.nih.gov/projects/msaviewer) to highlight conserved regions and polymorphisms.

To analyse variation in natural populations, short-read Illumina sequence data from 524 *P. falciparum* infections from nine different countries were aligned to a reference sequence from the *P. falciparum* IT line (PlasmoDB-68_PfalciparumIT_Genome.fasta, PIMMS43: PfIT_06: 789349-791016) using BWA MEM 0.7.17. This reference was used for sequence mapping as it has a full-length *PIMMS43* allele with no deletion (whereas the usual reference strain 3D7 has the large 150 bp deletion). The infections analysed each contained single predominant genotypes (as defined by within-infection fixation index *F*_WS_ values > 0.95 for each ^23,24^), selected from a larger database of available sequences to obtain a sample size of at least 50 from each of nine different countries covering the global endemic range of *P. falciparum*. These sequences were obtained from the Pf7 release of the global MalariaGEN data ^2^, from the following countries: Tanzania (N = 100), Malawi (N = 53), Democratic Republic of the Congo (N = 54), Cameroon (N = 53), Ghana (N = 51), The Gambia (N = 53), Colombia (N = 54), Cambodia (N = 53) and Papua New Guinea (N = 53).

Long indels were genotype-called from the aligned short-read sequence data using DELLY 1.7.2, leading to the identification of two different deletion variants (150 bp and 90 bp). SNP variants were called using GATK HaplotypeCaller 4.6.1.0. The variation in allele frequencies across the different populations was analysed using the *F*_ST_ index. Pairwise *F*_ST_ values for each of the polymorphisms between each population were estimated according to Weir and Cockerham’s θ index, using the R package hierfstat (https://github.com/jgx65/hierfstat) ^25^.

### PCR genotyping of *PIMMS43* indel polymorphisms

To amplify the central region of the coding sequence of *PIMMS43* using alternative nested PCR options, three pairs of oligonucleotide primers (Figure 2 and Supplementary Table 2) were designed to anneal to sequences conserved among all 20 *P. falciparum* strains aligned in initial analysis of variation (Figure 1). First round PCR with outer primers F1/R1 (Supplementary Table 2) was performed using an initial denaturation step at 95°C for 5 minutes, and 40 cycles at 95°C for 1 minute, 57°C for 45 seconds and 72°C for 2 minutes, followed by a final 5 minutes at 72°C. Products were diluted 1:100 with nuclease-free water and used as template for a second round nested PCR using primers F2/R2 or F3/R3 (Supplementary Table 2), using the same cycling conditions as for the first round PCR except for a shorter annealing time of 30 seconds at 60°C.

**Figure 1.**
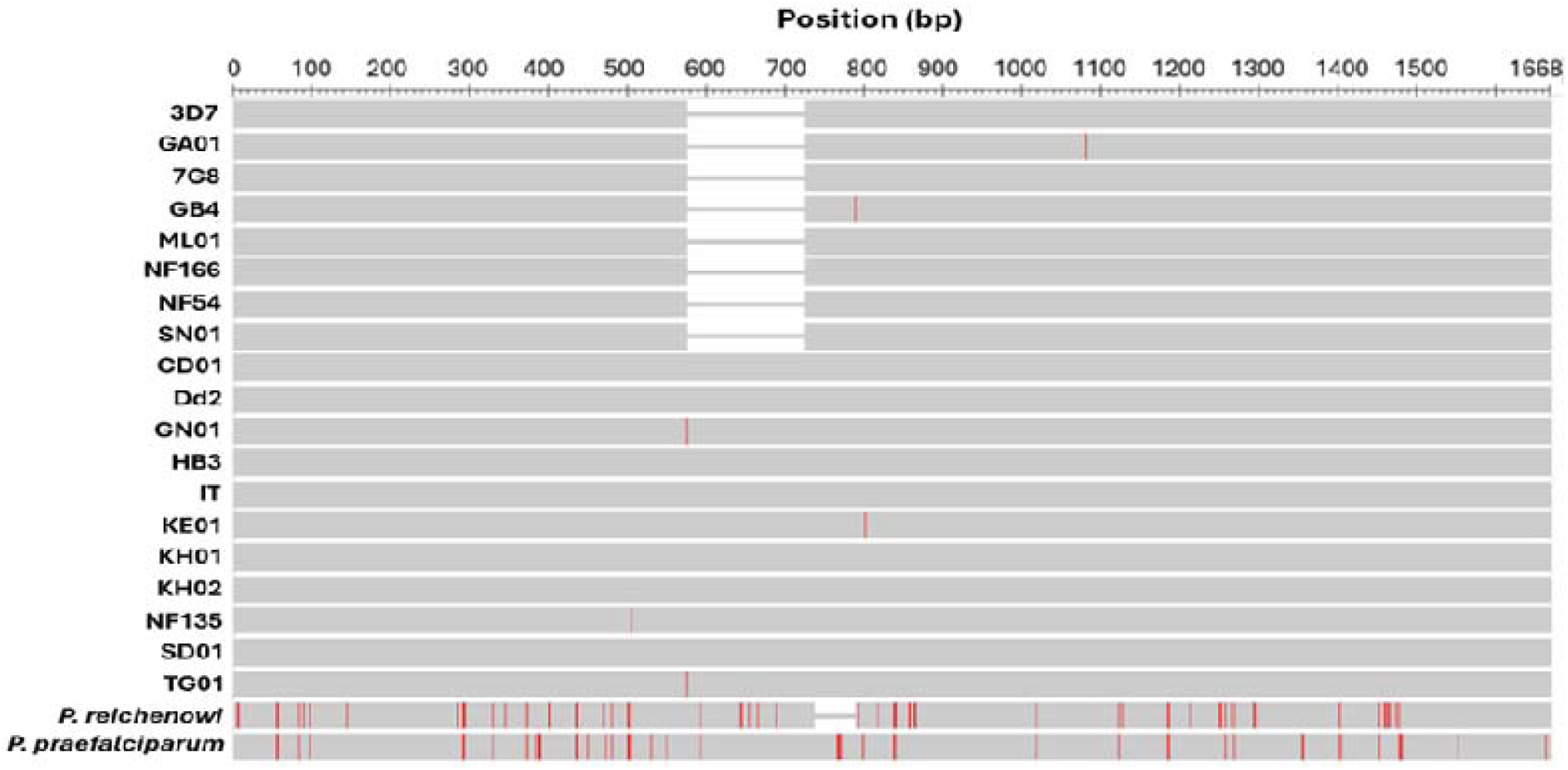
A large indel polymorphism in *P. falciparum PIMMS43* identified by comparison of diverse strains and closely-related species. Clustal alignment of *PIMMS43* coding sequences of diverse *P. falciparum* laboratory strains and clinical isolates with PacBio long-read genome sequence data ^3,4,12^, and individual isolates of the chimpanzee parasite *P. reichenowi* and the gorilla parasite *P. praefalciparum* ^20^. The scale covers the 1668 bp length of alleles without any deletion (including the laboratory strains Dd2 and IT). A 150 bp deletion allele in *P. falciparum* is seen at positions 573 – 722 in the top eight strains including 3D7 (the most widely-used *P. falciparum* reference) ^12^. Five SNPs are indicated with vertical red lines (at positions 505, 575, 787, 800 and 1081 numbered according to the intact Dd2/IT-like sequence). The *P. praefalciparum* orthologue has the same 1668 bp length as *P. falciparum* strains without the deletion while the *P. reichenowi* orthologue has a separate 54 bp deletion at positions 734 – 787. The sources and accession numbers of these sequences are given in Supplementary Table 1.

**Figure 2.**
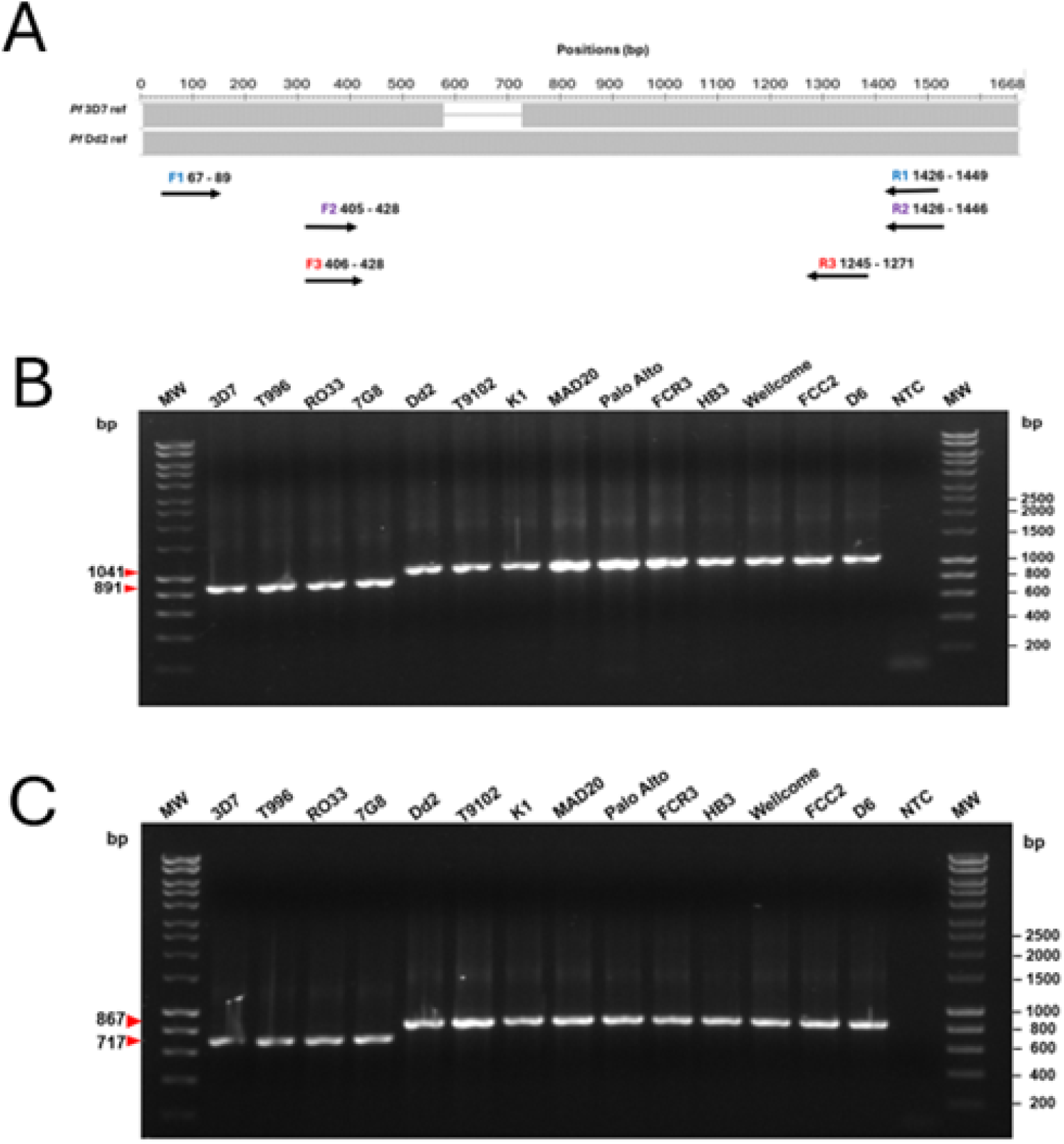
PCR detection of the *PIMMS43* 150 bp indel polymorphism among cultured *P. falciparum* lines. **A**. Illustration of primer design to capture the large indel (positions 573 to 722 of the Dd2 sequence missing in 3D7). Arrows indicate approximate positions of primer sequences (not to scale as oligonucleotides are shorter than the arrows on the scheme). First round PCR products are generated by outer primer pair F1/R1 (blue), and two nested PCR options are shown with inner primer pairs F2/R2 (blue) or F3/R3 (red) (details in Supplementary Table 2). **B**. Nested PCR products visualised for 14 *P. falciparum* laboratory strains with primer pair F2/R2 showing the *PIMMS43* deletion in the four strains on the left. The expected product sizes for the intact sequence and deletion are 1041 bp and 891 bp, respectively. **C**. Nested PCR products for the same *P. falciparum* strains with primer pair F3/R3 having expected product sizes of 867 or 717 bp. Both nested PCR assays enable visual scoring of the indel in all samples. Outer lanes on gels show molecular weight (MW) markers with the sizes of these shown on the right, and arrowheads on the left show the expected allelic sizes of PCR products. Blank lanes show a no-template control (NTC).

PCR products were electrophoresed on 1% agarose gels in 1x TAE buffer, with ethidium bromide staining to visualise on a Gel Doc XR+ Documentation System (Bio-Rad) using ImageLab v6.1 software. The larger (150 bp) and smaller (90 bp) deletion variants were scored by visual inspection of the bands on gel images. PCR products from laboratory strains were sequenced using di-deoxy chain termination Sanger sequencing after treatment with 3.33U Exonuclease I, 1U/uL FastAP Thermosensitive Alkaline Phosphatase, 1x FastAP buffer (ThermoFisher Scientific, UK) by Azenta Life Sciences (UK).

## Results

### Identification of a large deletion variant in the *P. falciparum PIMMS43* gene

To identify common polymorphisms within *PIMMS43*, coding sequences were compared among 19 *P. falciparum* lines for which PacBio long-read genome sequences were available ^3^. The most prominent and common polymorphism is a 150 bp indel (Figure 1), with eight of the *P. falciparum* lines having the deletion variant. This indel had not been previously noted, as analyses of *P. falciparum* polymorphisms normally have used the 3D7 genome sequence (which has the deletion) as a reference for mapping sequences. Apart from the indel only five single nucleotide polymorphisms (SNPs) were seen among the aligned *PIMMS43* allele sequences (Figure 1).

Comparison with the *PIMMS43* orthologue in the related species *P. praefalciparum* and *P. reichenowi* shows that both have a complete sequence, almost identical to the *P. falciparum* sequence without the deletion. The *P. praefalciparum* orthologue has the same 1668 bp coding sequence length as *P. falciparum* strains without the deletion, but there is a separate 54 bp deletion in the *P. reichenowi* sequence that begins 12 bp downstream from the position of the *P. falciparum* indel (Figure 1). As expected, there are multiple single nucleotide differences between each of these species and *P. falciparum* (including 18 positions where *P. praefalciparum* and *P. reichenowi* have the same nucleotide which is distinct from that in *P. falciparum*).

### Confirmation and genotyping of the large indel in *PIMMS43* by nested PCR

Identification of the 150 bp indel in *P. falciparum* encouraged an approach to confirm and further explore *PIMMS43* variation. First, the central region of the *PIMMS43* gene (primer sequences and positions shown in Supplementary Figure 2 and Supplementary Table 1) was amplified from a panel of 14 *P. falciparum* cultured laboratory strains. Use of alternative pairs of primers for nested PCR showed concordant expected product sizes, enabling straightforward visual scoring of the indel polymorphism (Figure 2). Sequencing of PCR products confirmed the indel with each *P. falciparum* line having either the Dd2-like (complete) or 3D7-like (150 bp deletion) sequence (Supplementary Figure 1). Apart from the indel, these sequences were highly conserved, with only four SNPs detected (of which two had been seen among the sequences illustrated in Figure 1). Combining with the sequences in Figure 1 identified 7 SNPs overall, with only one variant being seen in more than two of the strains (Supplementary Table 3).

### *PIMMS43* variants in population samples of human *P. falciparum* infections

To investigate the occurrence of the large deletion variant in natural populations and survey for additional deletion variants or common SNPs, *PIMMS43* sequences were inspected in genome sequences from population samples of malaria cases in nine countries, representing a wide range of the global endemic distribution. Six of the countries span the widespread distribution of *P. falciparum* within Africa. Analyses focused on infections that were genomically unmixed, with whole-genome short-read Illumina sequences in a global database ^2^ (the selected samples for this analysis had genome-wide *F*_WS_ values > 0.95 which indicates minimal within-infection diversity ^23^). Sequences from at least 50 such infections in each of the nine countries were mapped against the sequence from the IT strain as a reference, as this contains a complete *PIMMS43* sequence (Dd2-like) without any deletion (Figure 1).

This showed the 150 bp deletion variant to be common and widespread, and also identified a variant with a smaller (90 bp) deletion in the same region of the gene (Figure 3A). As neither of these deletions introduce a frameshift or nonsense codon, the variant alleles encode proteins missing 50 and 30 amino acids respectively, compared with full-length alleles and orthologues. The 150 bp deletion variant was detected in all African populations and in two of three non-African populations, whereas the small (90 bp) deletion was detected in most of the African populations but none of the non-African populations (Figure 3B and Table 1).

**Figure 3.**
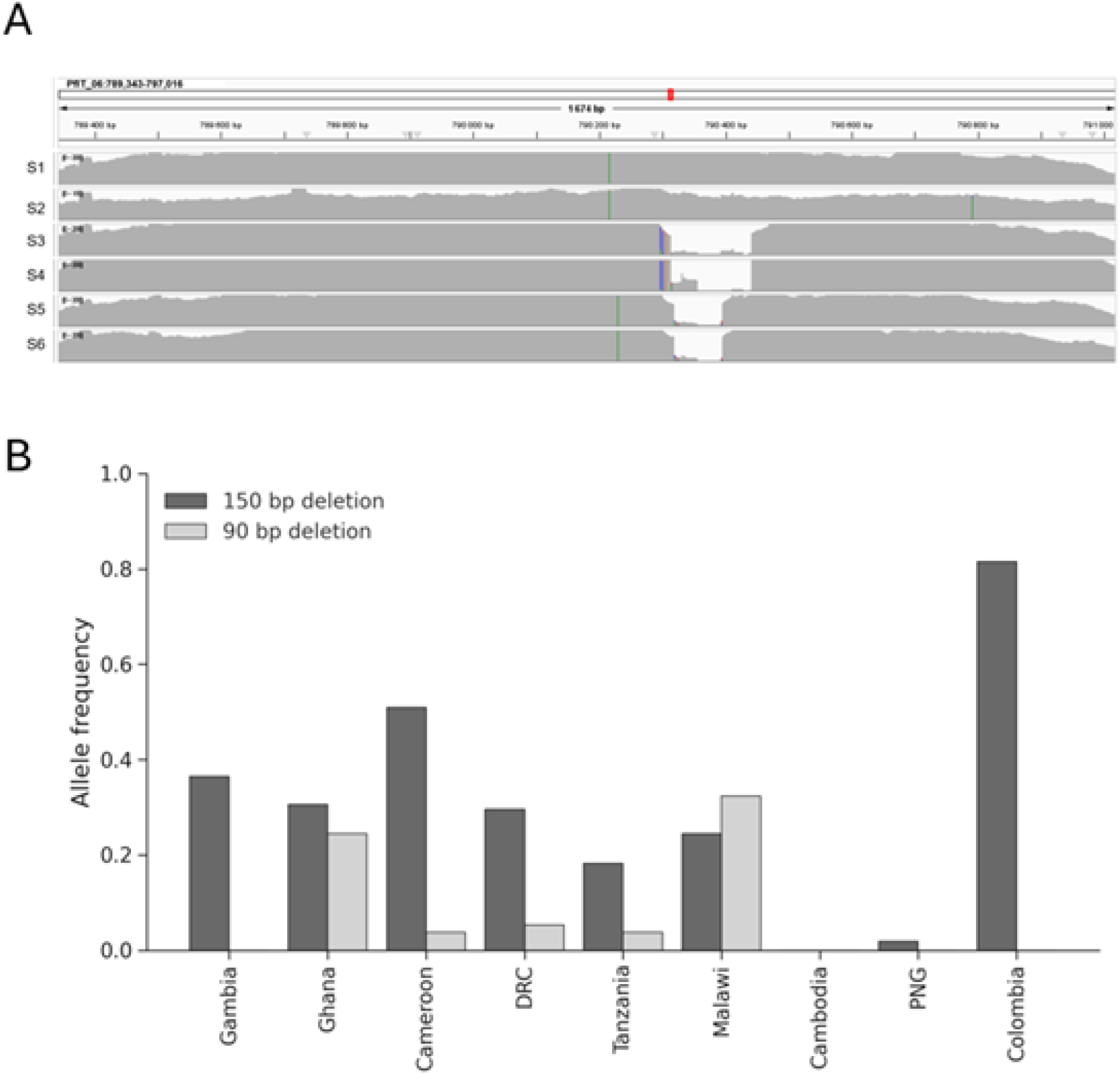
Two deletion variants in the *PIMMS43* gene are common in natural *P. falciparum* infections in endemic populations. **A**. Mapping of short-read Illumina sequences of genetically unmixed *P. falciparum* infections to a reference sequence from clone IT show examples with the large deletion variant (150 bp, infections S3 and S4) and a shorter deletion variant (90 bp, infections S5 and S6) as well as those with no deletion (infections S1 and S2). The screenshot from IGV (Ref) shows a few mismatches adjacent to the deletion as an artefact caused by the deletion. **B**. Frequency distribution of both deletion variants in nine countries (including six in Africa) with whole-genome short-read Illumina sequences in the MalariaGEN global database, focusing on infections that were genetically unmixed (N > 50 for each country with exact sample sizes listed in the Methods). The large (150 bp) deletion was detected at moderate frequencies in all African populations, and at more divergent frequencies in non-African populations. The small (90 bp) deletion variant was detected in most of the African populations, usually at a lower frequency than the large deletion.

**Table 1.**
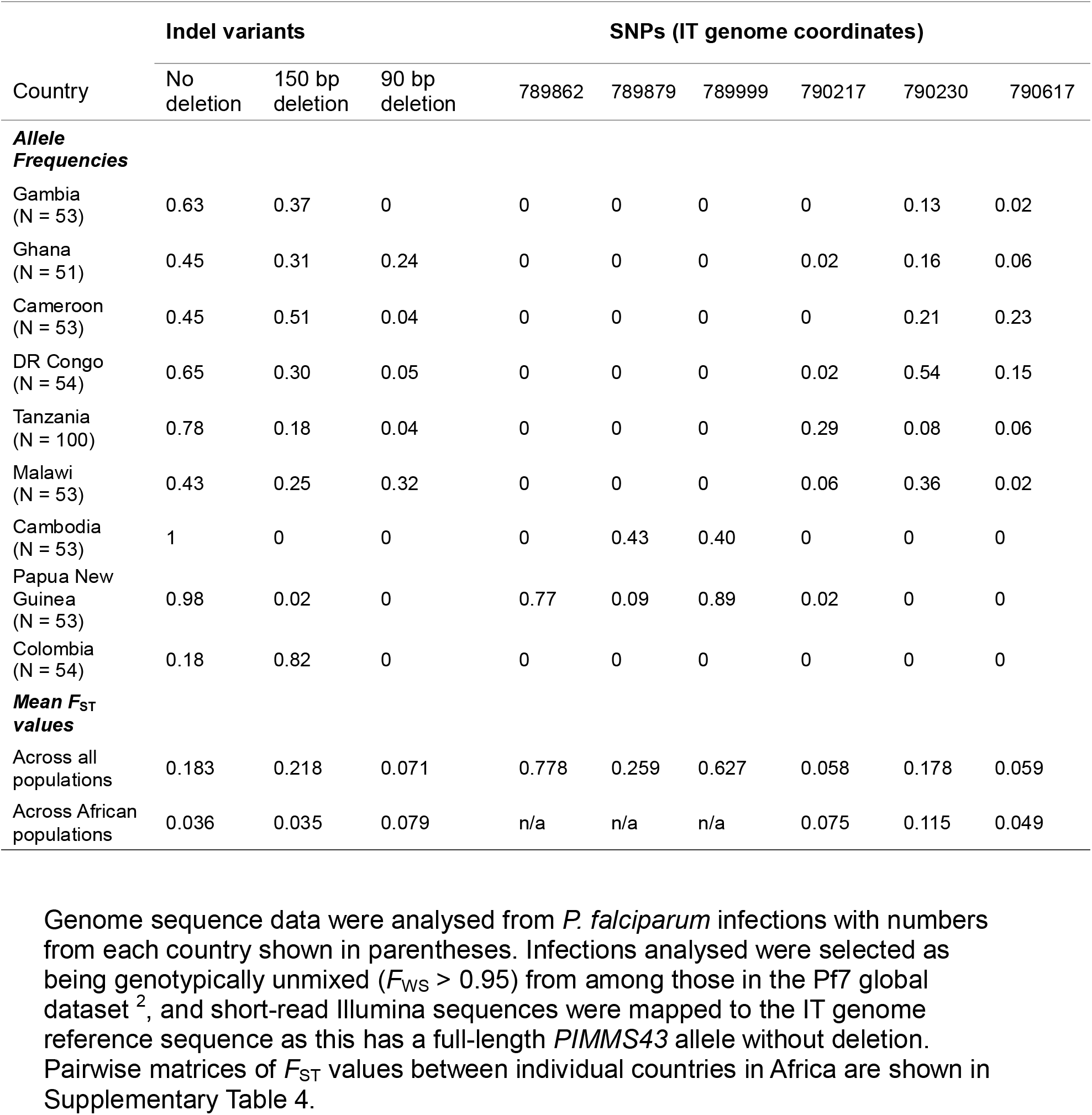
Variation in *PIMMS43* indel variant and SNP frequencies in *P. falciparum* population samples of human infections in endemic countries.

| Country | Indel variants |  |  | SNPs (IT genome coordinates) |  |  |  |  |  |
| --- | --- | --- | --- | --- | --- | --- | --- | --- | --- |
|  | No deletion | 150 bp deletion | 90 bp deletion | 789862 | 789879 | 789999 | 790217 | 790230 | 790617 |
| **Allele Frequencies** |  |  |  |  |  |  |  |  |  |
| Gambia (N = 53) | 0.63 | 0.37 | 0 | 0 | 0 | 0 | 0 | 0.13 | 0.02 |
| Ghana (N = 51) | 0.45 | 0.31 | 0.24 | 0 | 0 | 0 | 0.02 | 0.16 | 0.06 |
| Cameroon (N = 53) | 0.45 | 0.51 | 0.04 | 0 | 0 | 0 | 0 | 0.21 | 0.23 |
| DR Congo (N = 54) | 0.65 | 0.30 | 0.05 | 0 | 0 | 0 | 0.02 | 0.54 | 0.15 |
| Tanzania (N = 100) | 0.78 | 0.18 | 0.04 | 0 | 0 | 0 | 0.29 | 0.08 | 0.06 |
| Malawi (N = 53) | 0.43 | 0.25 | 0.32 | 0 | 0 | 0 | 0.06 | 0.36 | 0.02 |
| Cambodia (N = 53) | 1 | 0 | 0 | 0 | 0.43 | 0.40 | 0 | 0 | 0 |
| Papua New Guinea (N = 53) | 0.98 | 0.02 | 0 | 0.77 | 0.09 | 0.89 | 0.02 | 0 | 0 |
| Colombia (N = 54) | 0.18 | 0.82 | 0 | 0 | 0 | 0 | 0 | 0 | 0 |
| **Mean $F_{ST}$ values** |  |  |  |  |  |  |  |  |  |
| Across all populations | 0.183 | 0.218 | 0.071 | 0.778 | 0.259 | 0.627 | 0.058 | 0.178 | 0.059 |
| Across African populations | 0.036 | 0.035 | 0.079 | n/a | n/a | n/a | 0.075 | 0.115 | 0.049 |

The 150 bp deletion variant has a high *F*_ST_ index for variation when all populations are considered together, particularly due to having very divergent frequencies among non-African countries, being very common in Colombia and absent in Cambodia for example (Table 1). Across the African populations, the mean allele frequency of the 150 bp deletion variant was 31.7 % and the mean frequency of the 90 bp deletion variant was 11.6 %. The variation in deletion variant frequencies among African population samples is moderate, with frequencies of the 150 bp deletion ranging from 18 % in Tanzania to 51 % in Cameroon (mean *F*_ST_ of 0.035 among African populations), and frequencies of the 90 bp deletion ranging from zero in The Gambia to 32 % in Malawi (mean *F*_ST_ of 0.079 among African populations)

Only six SNPs within *PIMMS43* had minor allele frequencies of at least 5% overall (Table 1), and three of these had variants only in non-African populations. The polymorphisms with the most marked geographical divergence were those SNPs with variants only in non-African populations, having high variant frequencies only in Cambodia and Papua New Guinea which yielded high overall *F*_ST_ values (Table 1). Of the three SNPs with variants in Africa, the mean minor allele frequencies in African populations were 6.7% (for position 790217), 24.5 % (for position 790230), and 8.9 % (for position 790617). The variation in frequencies of these SNPs in Africa were moderate (Table 1 and Supplementary Table 4). It is notable that there was no SNP with a variant frequency as common as the 150 bp deletion, and only one SNP with a variant frequency as common as the 90 bp deletion (Table 1).

### *PIMMS43* indel variants in oocysts from wild-caught mosquitoes

To investigate the frequency of these major indel variants in naturally infected mosquitoes, PCR was performed on DNA isolated from individual oocysts (Figure 4). The central region of *PIMMS43* was successfully amplified from DNA samples from 30 individual *P. falciparum* oocysts from 25 wild-caught mosquitoes (11 *Anopheles gambiae s*.*s*. and 14 *An. funestus s*.*l*.) previously sampled in Tanzania ^15^. Each oocyst had a single predominant band corresponding to the expected size for one of the alleles, indicating all homozygote genotypes (Figure 4 and Table 2). Although only a few mosquitoes had more than one oocyst analysed, results show examples of different oocyst genotypes occurring together within infections in both *An. gambiae* and *A. funestus* (Supplementary Table 5).

**Figure 4.**
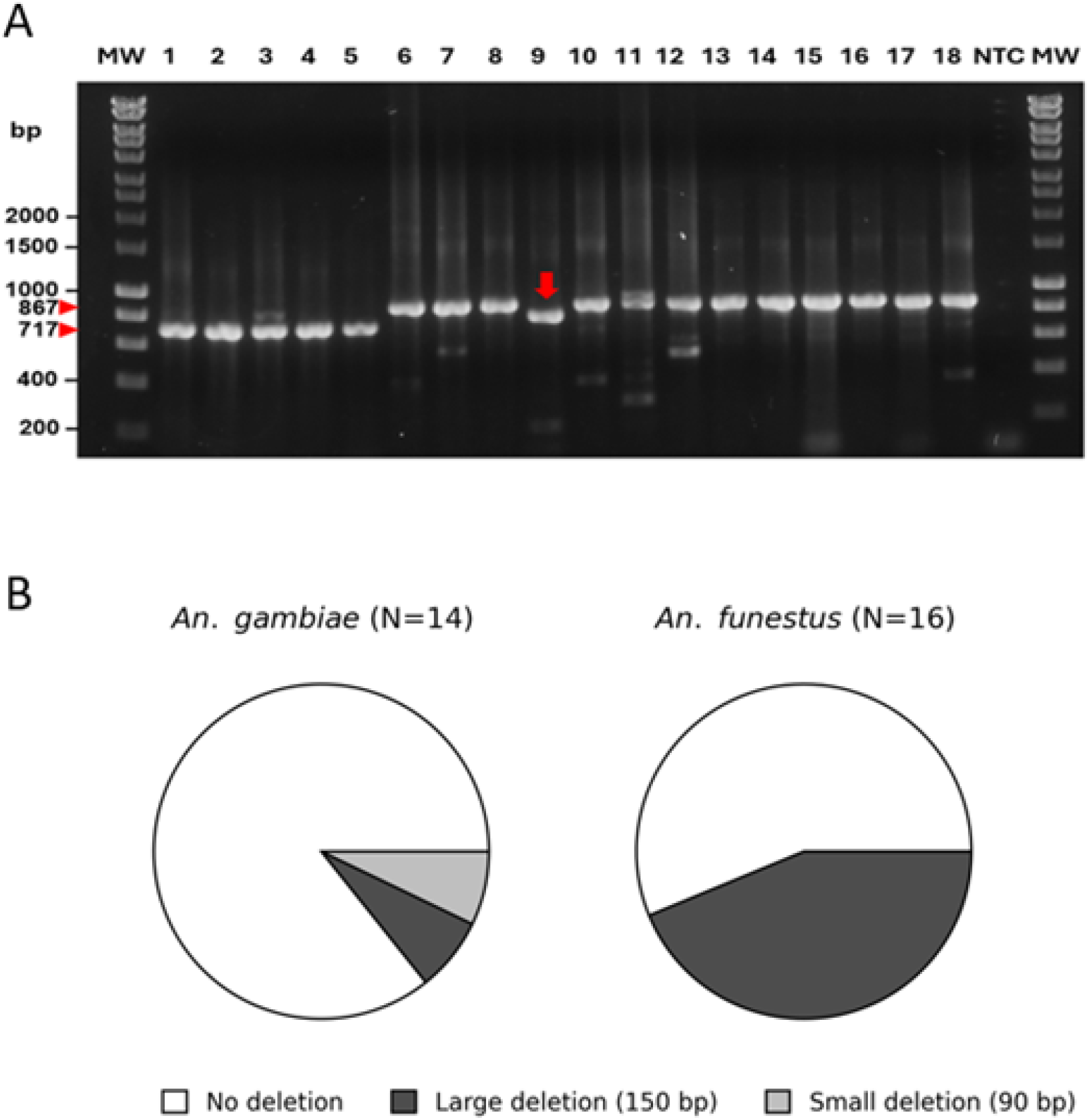
*PIMMS43* indel variants in *P. falciparum* oocysts from wild-caught mosquitoes. **A**. PCR products electrophoresed on a 1% agarose gel illustrate variant genotypes in 18 individual oocysts. Oocysts were isolated from *An. gambiae* and *An. funestus* mosquitoes in Tanzania and PCR amplification from extracted DNA used outer primers F1/R1 and nested primers F3/R3 (illustrated in Figure 2). Sizes of products corresponding to no-deletion (867 bp) and the 150 bp deletion (717 bp) are indicated in the side panel arrowheads, and the product of intermediate size corresponding the 90 bp deletion is marked with a red arrow. MW, 1 kb molecular weight marker; NTC, no template control. All oocysts shown have homozygote genotypes. **B**. Relative frequencies of the indel genotypes (all homozygotes) in oocysts according to the mosquito species from which each was isolated. Mosquitoes were sampled in the same area of Tanzania and the numbers of each genotype are shown in Table 1. There was a significant difference in *PIMMS43* indel genotype proportions in the sample of oocysts from the different species (Fisher’s Exact P = 0.039).

**Table 2.** *PIMMS43* indel polymorphisms in individual oocysts from mosquitoes in Tanzania.

| species | Numbers of oocysts from each mosquito |  |  |
| --- | --- | --- | --- |
|  | *An. gambiae*<br>(N = 14)<sup>a</sup> | *An. funestus*<br>(N = 16)<sup>b</sup> | Total<br>(N = 30) |
| ***PIMMS43* indel variant** |  |  |  |
| No deletion | 12 | 9 | 20 |
| Large deletion (150 bp) | 1 | 7 | 9 |
| Small deletion (90 bp) | 1 | 0 | 1 |

Across all 30 oocysts from the wild-caught mosquitoes, 20 had the intact no-deletion genotype, 9 had the larger (150 bp) deletion, and one had the smaller (90 bp) deletion (Figure 4). Overall, the *PIMMS43* indel allele frequencies among the Tanzanian oocysts (0.67 for the no-deletion type, 0.30 for the 150 bp deletion, and 0.03 for the 90 bp deletion) were similar to the frequencies seen in the human infections in Tanzania (Figure 3 and Table 1).

A comparison of the relative proportions of the different *PIMMS43* variants in oocysts from the two mosquito species indicates a difference of borderline statistical significance. Analysis of all oocysts indicates a significant skew in genotype proportions between the mosquito species (Fisher’s Exact P = 0.039) (Figure 4B and Table 2). However, analysis of only one oocyst from each mosquito (by random sampling to exclude data from additional oocysts) reduced the significance of the difference (Fisher’s Exact P = 0.075) (Table 2).

## Discussion

Large deletion variants are described in a functionally important gene for malaria parasite transmission, and their frequencies in human infections different populations are determined, alongside samples from vector mosquitoes. The discovery of these variants illustrates the value of having diverse isolates characterised by long-read sequencing. In this case, a multiple alignment of the sequences of different isolates previously obtained ^3,4^ revealed the large (150 bp) deletion in several clinical isolates and laboratory strains, including the usual genome reference strain 3D7. Choosing a strain that had the intact no-deletion allelic sequence as a reference allowed mapping of short-read sequences from natural infections, showing the widespread distribution of this large deletion as well as an alternative smaller (90 bp) deletion in the same region of the gene.

The identification of these variants in *PIMMS43* suggests that deletion variants may remain to be discovered in many other genes, as structural variation has not been as fully characterised as SNP variation in large global surveys of genome sequences from natural infections ^1,2^. Some undiscovered structural variants are likely to contribute to parasite adaptation, and be targets of strong selection in natural populations, as has been illustrated for copy number variation ^26-28^. As most population genetic studies of malaria parasites have analysed SNP variation, identification of loci has usually oriented around SNPs for analyses of temporal selection ^29^, geographically-varying selection ^30^, or associations of parasite and host genotypes ^31,32^. However, in many cases the signatures of selection are due to SNPs in tight linkage disequilibrium with causal variants that remain unidentified, some of which may be large indels or other structural variants.

Further long-read sequencing of a larger sample of *P. falciparum* clinical isolates would be required to survey the frequency and distributions of large indels and other structural variation throughout the genome. Recently derived long-read sequences of a collection of isolates from one endemic area ^33^ shows that such data may be forthcoming to enable this. Once the common naturally occurring indels are clearly described, it should be possible to genotype these with short-read sequencing, mapping to reference sequences that contain representative full-length alleles. Ideally, constructing pangenome reference sequences would facilitate this process, as has been done for human polymorphism analysis ^34^.

A previous study that demonstrated the functional importance of *PIMMS43* presented an exploratory analysis of single nucleotide polymorphism in the gene in different African countries ^8^. Although the allele frequencies were not presented, it was suggested that SNPs had higher *F*_ST_ scores than might be expected in comparisons between West Africa to East or Central Africa. Here it is shown that most SNPs in *PIMMS43* have very low overall minor allele frequencies, so we analysed the frequency distributions of SNPs that had overall minor allele frequencies of at least 5 %. Only three such SNPs are seen in *PIMMS43* in African *P. falciparum* populations, and these showed moderate variation in frequencies among African populations, similar to the level of geographic variation in the indel frequencies.

The similar indel allele frequencies observed in human infections in Tanzania and in oocysts from wild-caught mosquitoes is as expected, given there is no previous evidence of major geographical population structure or subdivision of *P. falciparum* within Tanzania over the time when the samples were collected. More recently, as malaria has declined in parts of Tanzania, minor differences in parasite population genetic composition are becoming apparent in different parts of the country ^35^. The difference in the frequencies of the PIMMS43 variants between the mosquito species, although based on a small sample size, highlights the importance of conducting larger surveys of parasite genetic variation in different mosquito vectors. Such expanded genomic analyses of parasites within mosquito vectors will contribute to a more complete understanding of selection on parasite populations.

In this analysis, with a limited sample size of oocysts from wild-caught mosquitoes, only homozygotes for the indel polymorphism were seen. This is not unexpected given parasite inbreeding that occurs due to self-fertilisation in infections containing a limited pool of gametes, as previously estimated in this population and elsewhere by analysis of microsatellite markers and other polymorphisms ^13,15,16,36^. Genotyping larger sample sizes of oocysts should enable analysis of heterozygotes and potential variation in inbreeding coefficients among parasite loci, which may detect hidden targets of selection ^15,16^. Obtaining larger sample sizes of individual oocysts from natural infections is difficult due to a generally very low infection prevalence in mosquitoes and technical challenge in isolating individual oocysts. Analysis of parasite genotypes within DNA extractions from whole mosquito abdomen or head samples may be another means of surveying more infections ^37^.

Alternatively, as larger numbers of oocysts may be obtainable from experimental infections of colony-reared mosquitoes, parasite genotype distributions could be studied in such infections ^38^, and potentially compared between colonies of different mosquito species. A previous discovery of selection on the *Pfs47* gene ^15^ that encodes another ookinete protein ^39^, encouraged a long series of experimental studies that revealed a mechanism of allelic evasion of innate mosquito immunity ^40,41^. If further studies implicate the *PIMMS43* variants as being under selection, experimental studies on their phenotypic effects will be warranted. Moreover, it is likely that large indel variants in other genes will be discovered by future analyses of long-read genome sequences, and the phenotypic effects of some of these will need investigation.

## Ethical Approval

This study was submitted to the Ethics Committee of the London School of Hygiene and Tropical Medicine and approved as being exempt from requiring further review, as all parasite genomic data analysed from human infections were already in the public domain as part of the MalariaGEN PF7 data release (https://www.malariagen.net/resource/34/), and parasite genotyping was on samples from laboratory strains and mosquitoes that did not include human subjects or require ethical approval for genotyping.

