## Supplementary for "Large deletion variants in a *Plasmodium falciparum* ookinete protein gene and associations with different endemic populations and mosquito vectors"

**Supplementary Information**

**Supplementary Table 1.** Gene IDs of *PIMMS43* sequences from different *P. falciparum* lines and related species with long-read genome sequences used for the alignment in Figure 1.

| Species/Strain | Gene ID [a] | *PIMMS43* indel type |
|---|---|---|
| *P. falciparum* 3D7 | Pf3D7_0620000 | 150 bp deletion |
| *P. falciparum* GA01 | PfGA01_060025000 | 150 bp deletion |
| *P. falciparum* 7G8 | Pf7G8_060024900 | 150 bp deletion |
| *P. falciparum* 7G8-2 | Pf7G8-2_000172300 | 150 bp deletion |
| *P. falciparum* GB4 | PfGB401_060024400 | 150 bp deletion |
| *P. falciparum* ML01 | PfML01_060023800 | 150 bp deletion |
| *P. falciparum* NF166 | PfNF166_06002440 | 150 bp deletion |
| *P. falciparum* NF54 | PfNF54_060025000 | 150 bp deletion |
| *P. falciparum* SN01 | PfSN01_060025300 | 150 bp deletion |
| *P. falciparum* CD01 | PfCD01_060025400 | no deletion |
| *P. falciparum* Dd2 | PfDd2_060024700 | no deletion |
| *P. falciparum* GN01 | PfGN01_060025700 | no deletion |
| *P. falciparum* HB3 | PfHB3_060024200 | no deletion |
| *P. falciparum* IT | PfIT_060023800 | no deletion |
| *P. falciparum* KE01 | PfKE01_060026100 | no deletion |
| *P. falciparum* KH01 | PfKH01_060027000 | no deletion |
| *P. falciparum* KH02 | PfKH02_060026500 | no deletion |
| *P. falciparum* NF135 | PfNF135_060024200 | no deletion |
| *P. falciparum* SD01 | PfSD01_060023900 | no deletion |
| *P. falciparum* TG01 | PfTG01_060025300 | no deletion |
| *P. reichenowi* | PRCDC_0618400 | 54 bp deletion |
| *P .praefalciparum* | PPRFG01_0618500 | no deletion |

[a] Gene sequences obtained from PacBio sequences of different isolates (published data from Otto et al. 2018a https://doi.org/10.12688/wellcomeopenres.14571.1, Otto et al 2018b https://doi.org/10.1038/s41564-018-0162-2, Moser et al. 2020 https://doi.org/10.1186/s13073-019-0708-9) and retrieved using PlasmoDB (www.plasmodb.org).

**Supplementary Table 2.** Primers used for PCR amplification of *P. falciparum PIMMS43* to analyse polymorphisms in the central part of the coding sequence (in Figure 2 and Supplementary Figure 1).

| Primer pairs | Expected product size (bp) | |
|---|---|---|
| | 150 bp deletion | No deletion |
| F1 – 5' CTTTCCAAAGATGATGGCCAACG 3' | 1233 | 1383 |
| R1 – 5' TGAGCATTCATGGAGAAAATCGTC 3' | | |
| F2 – 5' GAGGAAGATATGTTTGTAGAAGG 3' | 891 | 1041 |
| R2 – 5' GCATTCATGGAGAAAATCGTC 3' | | |
| F3 – 5' TGAGGAAGATATGTTTGTAGAAGG 3' | 717 | 867 |
| R3 – 5' GAACTAATCGAATGATTATATTTAACC 3' | | |

A product size difference of 150 bp is expected depending on whether alleles have the following sequence deleted (5'*AAAGGAAATTGAAGATGAAGGAAGAAGGAGAAAAGATATTGAAGAAGAAAATAGAAAGAAAAAGGAATTAAAGGATGCCCAAAGAAAGAAGGAAGATATGGAATATCAAGAAATAATGAAAAAATATATTGAAGAAGAAGAAAAAATGAA* 3').

**Supplementary Table 3.** *PIMMS43* SNPs identified in *P. falciparum* sequences aligned in Figure 1 and Supplementary Figure 2**.**

| SNP position in full-length (no del) sequence | Reference Allele | Alternate allele | Amino acid | *P. falciparum* strains with alternate allele |
|---|---|---|---|---|
| 505 [a,b] | C | A | Q169K | NF153, T9102 |
| 575 [a,b] | A | G | K192R | D6, GN01, TG01 |
| 787 | G | T | D213Y | GB4 |
| 800 [a] | C | T | S267L | KE01 |
| 966 [b] | T | A | I322I (syn) | D6 |
| 1018 [b] | G | A | D290N | MAD20, K1 |
| 1081 [a] | C | A | L311I | GA01 |

[a] SNPs identified among reference sequences in Figure 1.
[b] SNPs identified among laboratory strains in Supplementary Figure 1.

**Supplementary Table 4.** Matrices of pairwise $F_{ST}$ values between each population in Africa for each of the PIMMS43 polymorphisms. Higher values showing the most pairwise divergence are highlighted by pink and red shading.

**150 bp deletion**

| | Cameroon | Gambia | Ghana | Malawi | RDC | Tanzania |
|---|---|---|---|---|---|---|
| Cameroon | | | | | | |
| Gambia | 0.03 | | | | | |
| Ghana | 0.06 | -0.02 | | | | |
| Malawi | 0.12 | 0.01 | -0.01 | | | |
| RDC | 0.07 | -0.01 | -0.02 | -0.01 | | |
| Tanzania | 0.20 | 0.06 | 0.03 | -0.01 | 0.02 | |

**90 bp deletion**

| | Cameroon | Gambia | Ghana | Malawi | RDC | Tanzania |
|---|---|---|---|---|---|---|
| Cameroon | | | | | | |
| Gambia | 0.00 | | | | | |
| Ghana | -0.03 | 0.00 | | | | |
| Malawi | 0.23 | 0.30 | 0.26 | | | |
| RDC | -0.03 | 0.02 | -0.02 | 0.20 | | |
| Tanzania | -0.02 | 0.01 | -0.02 | 0.31 | -0.02 | |

**No deletion allele**

| | Cameroon | Gambia | Ghana | Malawi | RDC | Tanzania |
|---|---|---|---|---|---|---|
| Cameroon | | | | | | |
| Gambia | 0.04 | | | | | |
| Ghana | 0.06 | -0.02 | | | | |
| Malawi | -0.02 | 0.02 | 0.03 | | | |
| RDC | 0.06 | -0.02 | -0.02 | 0.03 | | |
| Tanzania | 0.18 | 0.03 | 0.02 | 0.14 | 0.02 | |

**150bp / 90 bp / no del three allelic system**

| | Cameroon | Gambia | Ghana | Malawi | RDC | Tanzania |
|---|---|---|---|---|---|---|
| Cameroon | | | | | | |
| Gambia | 0.03 | | | | | |
| Ghana | 0.05 | -0.02 | | | | |
| Malawi | 0.08 | 0.06 | 0.05 | | | |
| RDC | 0.06 | -0.01 | -0.02 | 0.05 | | |
| Tanzania | 0.18 | 0.05 | 0.02 | 0.11 | 0.02 | |

**PfIT_06:790217 SNP**

| | Cameroon | Gambia | Ghana | Malawi | RDC | Tanzania |
|---|---|---|---|---|---|---|
| Cameroon | | | | | | |
| Gambia | NA | | | | | |
| Ghana | 0.00 | 0.00 | | | | |
| Malawi | 0.04 | 0.04 | 0.00 | | | |
| RDC | 0.00 | 0.00 | -0.02 | 0.00 | | |
| Tanzania | 0.23 | 0.23 | 0.19 | 0.14 | 0.20 | |

**PfIT_06:790230 SNP**

| | Cameroon | Gambia | Ghana | Malawi | RDC | Tanzania |
|---|---|---|---|---|---|---|
| Cameroon | | | | | | |
| Gambia | 0.00 | | | | | |
| Ghana | -0.01 | -0.02 | | | | |
| Malawi | 0.04 | 0.11 | 0.08 | | | |
| RDC | 0.19 | 0.30 | 0.26 | 0.04 | | |
| Tanzania | 0.06 | 0.00 | 0.02 | 0.22 | 0.43 | |

**PfIT_06:790617 SNP**

| | Cameroon | Gambia | Ghana | Malawi | RDC | Tanzania |
|---|---|---|---|---|---|---|
| Cameroon | | | | | | |
| Gambia | 0.17 | | | | | |
| Ghana | 0.09 | 0.00 | | | | |
| Malawi | 0.17 | -0.02 | 0.00 | | | |
| RDC | 0.00 | 0.09 | 0.02 | 0.09 | | |
| Tanzania | 0.11 | 0.00 | -0.02 | 0.00 | 0.03 | |

**Supplementary Table 5.** *PIMMS43* indel genotypes in 30 individual *P. falciparum* oocysts from wild-caught Tanzanian mosquitoes.

| Mosquito species | Mosquito ID | Oocyst ID | Indel type (all homozygotes) |
|---|---|---|---|
| *An. funestus* | 36-11 | 40 | No deletion |
| | 72-12 | 71* | No deletion |
| | 72-12 | 72* | 150bp del |
| | 72-12 | 73* | 150bp del |
| | 76-36 | 74 | 150bp del |
| | 72-19 | 87 | 150bp del |
| | 36-11 | 88 | No deletion |
| | 76-35 | 114 | 150bp del |
| | 36-1 | 170 | No deletion |
| | 95-1 | 214 | No deletion |
| | 95-2 | 217 | No deletion |
| | 73-2 | 221 | No deletion |
| | 95-3 | 347 | No deletion |
| | 92-9 | 356 | 150bp del |
| | 89-2 | 367 | 150bp del |
| | 83-17 | 368 | No deletion |
| *An. gambiae* | 36-3 | 27 | No deletion |
| | 46-1 | 46* | 90bp del |
| | 46-1 | 47* | No deletion |
| | 84-1 | 90 | No deletion |
| | 77-2 | 86 | No deletion |
| | 83-18 | 96 | No deletion |
| | 76-22 | 130 | No deletion |
| | 83-3 | 244 | No deletion |
| | 95-7 | 248 | No deletion |
| | 76-5 | 275 | 150bp del |
| | 83-16 | 285* | No deletion |
| | 83-16 | 286* | No deletion |
| | 89-12 | 288* | No deletion |
| | 89-12 | 289* | No deletion |

16 oocysts were from *An. funestus* s.l. mosquitoes (14 mosquitoes of which 3 oocysts were sampled from one), and 14 oocysts were from *An. gambiae* s.s. mosquitoes (11 mosquitoes, with 2 oocysts sampled from each of three). Asterisks show oocysts that are among more than one from an individual mosquito. Results show examples of different oocyst genotypes occurring within individual infections in both *An. gambiae* and *An. funestus*.

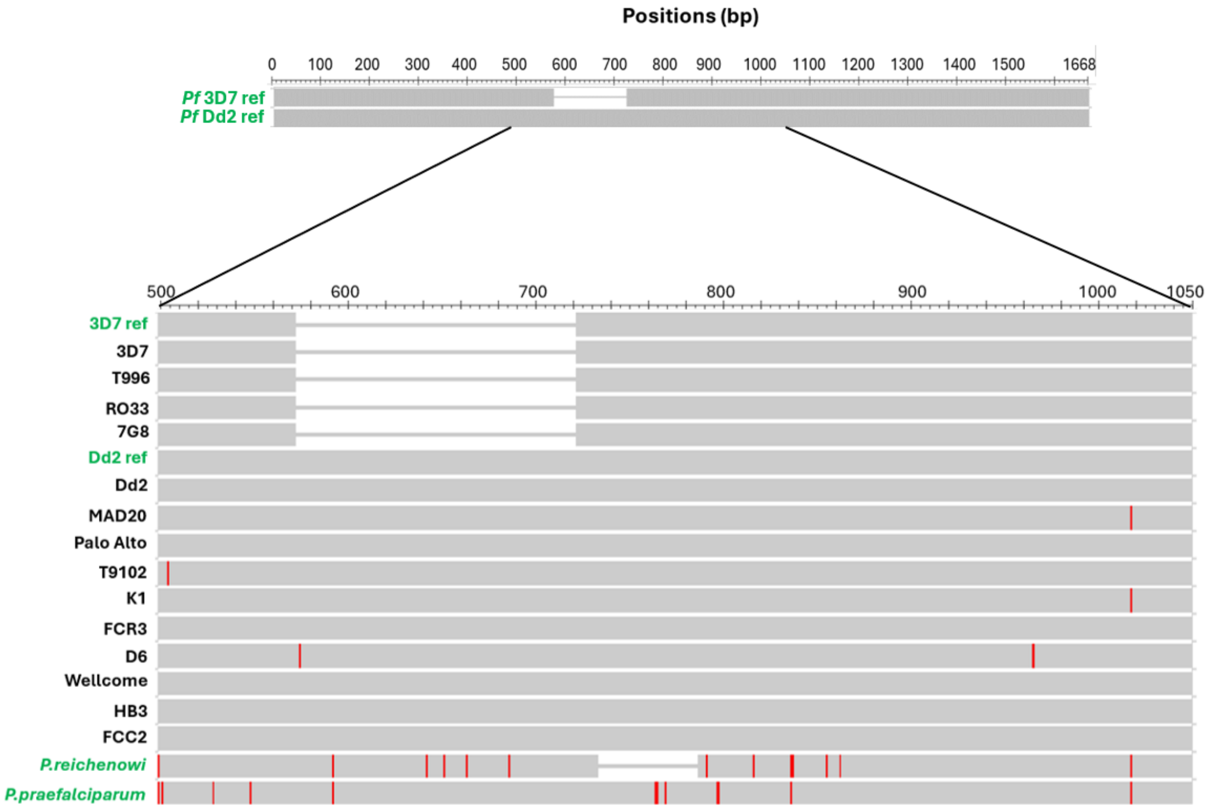


**Supplementary Figure 1. Confirmation of the *PIMMS43* large indel (150 bp) polymorphism among cultured *P. falciparum* laboratory strains.** Alignment of sequences of *PIMMS43* PCR products from 14 *P. falciparum* laboratory lines focusing on base pairs 500 – 1050 incorporating the indel and flanking regions. Conserved regions are shown as grey bars, deletions are indicated by gaps between bars. SNPs are indicated with vertical red lines showing differences from 3D7. Published sequences are indicated in green text (for 3D7, Dd2, *P. reichenowi* and *P. praefalciparum*). Three non-synonymous SNPs were observed at positions 505 (C → A), 575 (A → G) and 1018 (G→A), and one synonymous SNP was observed at position 966 (T → A). The SNPs at positions 505 and 575 had also been observed among the previously published sequences aligned in Figure 1.
